# Species tree-aware simultaneous reconstruction of gene and domain evolution

**DOI:** 10.1101/336453

**Authors:** Sayyed Auwn Muhammad, Bengt Sennblad, Jens Lagergren

## Abstract

Most genes are composed of multiple domains, with a common evolutionary history, that typically perform a specific function in the resulting protein. As witnessed by many studies of key gene families, it is important to understand how domains have been duplicated, lost, transferred between genes, and rearranged. Analogously to the case of evolutionary events affecting entire genes, these domain events have large consequences for phylogenetic reconstruction and, in addition, they create considerable obstacles for gene sequence alignment algorithms, a prerequisite for phylogenetic reconstruction.

We introduce the DomainDLRS model, a hierarchical, generative probabilistic model containing three levels corresponding to species, genes, and domains, respectively. From a dated species tree, a gene tree is generated according to the DL model, which is a birth-death model generalized to occur in a dated tree. Then, from the dated gene tree, a pre-specified number of dated domain trees are generated using the DL model and the molecular clock is relaxed, effectively converting edge times to edge lengths. Finally, for each domain tree and its lengths, domain sequences are generated for the leaves based on a selected model of sequence evolution.

For this model, we present a MCMC-based inference framework called DomainDLRS that takes a dated species tree together with a multiple sequence alignment for each domain family as input and outputs an estimated posterior distribution over reconciled gene and domain trees. By requiring aligned domains rather than genes, our framework evades the problem of aligning full-length genes that have been exposed to domain duplications, in particular non-tandem domain duplications. We show that DomainDLRS performs better than MrBayes on synthetic data and that it outperforms MrBayes on biological data. We analyse several zincfinger genes and show that most domain duplications have been tandem duplications, some involving two or more domains, but non-tandem duplications have also been common.

## 1 Introduction

The main evolutionary events that affects gene evolution include speciation, gene duplication, gene loss, incomplete lineage sorting, and lateral gene transfer. During the last 10-15 years, considerable attention has been given to how such events induce an interplay between gene and species evolution and, in particular, its consequences for phylogenetic reconstruction. This trend has inspired considerable method development culminating in probabilistic species tree-aware methods for gene tree reconstruction and methods for simultaneous reconstruction of gene trees and species trees [1, 2, 3]. Complicating the issue further, most genes are composed of multiple domains, each a segment of contiguous nucleotides with a common evolutionary history that typically performs a specific function in the resulting protein (although also structure-based definitions of domains are common). As shown by many studies of key gene families such as PRDM9, ZNF91, and Reelin [4, 5, 6], domains can be also be an appropriate organizational units in evolutionary analyses, e.g., it is important to understand how they have been duplicated, lost, transferred between genes, and rearranged. Similar to the case of evolutionary events affecting entire genes, these domain events have large consequences for phylogenetic reconstruction. In addition, they create considerable obstacles for gene sequence alignment algorithms, which constitute a prerequisite for phylogenetic reconstruction. Since domains are key functional units, an improved capacity to reconstruct the evolutionary history of domains and relating it to that of the hosting genes as well as species, will facilitate increasingly advanced evolutionary studies of gene and domain function, in particular by correlating evolutionary changes with metabolic, physiological, and morphological changes. Unfortunately, in contrast to methods for analyzing gene evolution, major advancements of methods for domain evolution studies have been scarce.

The majority of all proteins (66% in unicelluar organisms and more than 80% in metazoa) are multi-domain proteins [7] that have particular evolutionary as well as functional importance. These proteins perform many important functions including binding, catalytic, and signaling activities, making domains an essential concept also in construction and studies of regulatory, structural, and signaling networks. One interesting example is the zinc-finger gene family C2H2-ZNF, which mostly encodes DNA binding proteins that regulate gene expression by binding to DNA via an array of zinc finger domains [8]. This is the largest gene family among the human transcription factors, in fact constituting approximately a quarter of those.

The KRAB domain-containing C2H2-ZNF subfamily, KRAB-ZNF, which with a few exceptions represses the expression of target genes, has a striking number of lineagespecific expansions as well as domain duplications among primates [9], suggesting a major role for KRAB-ZNF family in regulatory evolution. Nowick et al [10] identified KRAB-ZNF genes as markedly over-represented among genes with significant differential expression in human and chimpanzee brain, thereby implicating the KRAB-ZNF family in human brain evolution. Considerable interest has also been devoted to the role of KRAB-ZNF genes in suppressing endogenous retroelements, i.e., endogenous retroviruses and nonretroviral retrotransposons [11]. This aspect of KRAB-ZNF functionality is statistically supported by a correlation between the emergence of new endogenous LTR (Long Terminal Repeat) elements and tandem duplications of ZNF genes [12]. The identity of the ZNF genes suppressing endogenous retroelements is largely unknown, but a recent study revealed that ZNF91 and ZNF93 from the KRAB-ZNF family have evolved to repress SINE-VNTR-Alus and long interspersed nuclear elements, which are two currently active human endogenous retrotransposons [5]. Although these evolutionary scenarios may appear entirely unrelated, they have been connected, for instance by the hypothesis that many KRAB-ZNFs evolved as repressors of endogenous retroelements, but was later been co-opted by the regulatory machinery and regulate genes in regions that used to host endogenous retroelements [12].

Goodman *et al.* [13] pioneered gene-species tree comparisons by introducing the *most parsimonious reconciliation* (MPR). MPR maps each gene tree vertex to either a species tree vertex, in which case it represents a speciation, or to a species tree edge, in which case it represents a duplication, in a way that minimizes the number of duplications. This classical paper by Goodman *et al.* on reconciling gene trees and species trees can be seen as the starting point for the development of models and methods capturing gene and domain evolution. Goodman *et al.* introduced reconciliations in order to explain how a gene family, represented by a gene tree, has evolved through gene duplications and losses relative to the associated species tree. Later extensions of this line of work gave rise to parsimony methods allowing also lateral gene transfers [14]. There are today also MPR methods directly targeting domain evolution. The straightforward domain architecture (DA) models have mainly been used to study domain distributions across gene and genomes. Many of the parsimony-based tools for gene and species evolution may equally well be used to study domain evolution relative to that of the associated gene family. In a recent effort to extend parsimony-based methods to also be applicable to domain analyses, Stolzer *et al.* introduce a method that reconciles a domain tree with a gene tree that has previously been reconciled with a species tree, under either the duplication-loss or duplication-transfer-loss model [15].

Recently, also integrated probabilistic models have been proposed for gene evolution combining sequence evolution under a relaxed molecular clock with gene duplication and loss, or even gene duplication, gene loss and lateral gene transfer [16, 1, 17]. Building on these integrated probabilistic models, so-called species tree-aware gene tree reconstruction methods that, apart from gene sequences, take advantage of a species tree, have been proposed and shown to perform superiorly [1, 2] to earlier methods. Åkerborg et al. [1] presented an MCMC-based Bayesian analysis framework, DLRS, that can be used to estimate the posterior distribution over gene trees. More precisely, the DLRS model incorporates submodels for gene tree evolution, sequence evolution and a relaxed molecular clock to define a posterior over gene trees given a species tree and the observed multiple sequence alignment. These approaches can, in addition to from being used to obtain more reliable gene trees and reconciliations, also incorporate what traditionally have been more down-stream analysis in order to obtain, e.g., improved orthology analysis methods, that takes advantage of the gene sequences directly, i.e., not merely when constructing the gene tree [18, 19]. Here we extend the DLRS model to also capture domain evolution.

## 2 The DomainDLRS Model

The DomainDLRS model is a hierarchical, generative probabilistic model containing three levels corresponding to species, genes, and domains, respectively. As the model is described below, a dated species tree *T*_*S*_, *t*_*S*_ is given rather than generated by the model, but it is easy to extend this version so also a species tree is generated, e.g., using a birthdeath process. The model is detailed below, but can briefly be described as follows. From a dated species tree, a gene tree *T*_*G*_ is generated according to the DL model; this also induces a dating of the vertices of the gene tree. Then, separate independent applications of the DL model is used to generate a pre-specified number of dated domain trees from the dated gene tree. For each domain tree, the R model is used to relax the molecular clock and effectively converts edge times to edge length. Finally, for each domain tree and its lengths domain sequences are generated for the leaves of based on a model of sequence evolution M (i.e., the user’s model of choice, any standard model of sequence evolution can be used).

**Figure 1:**
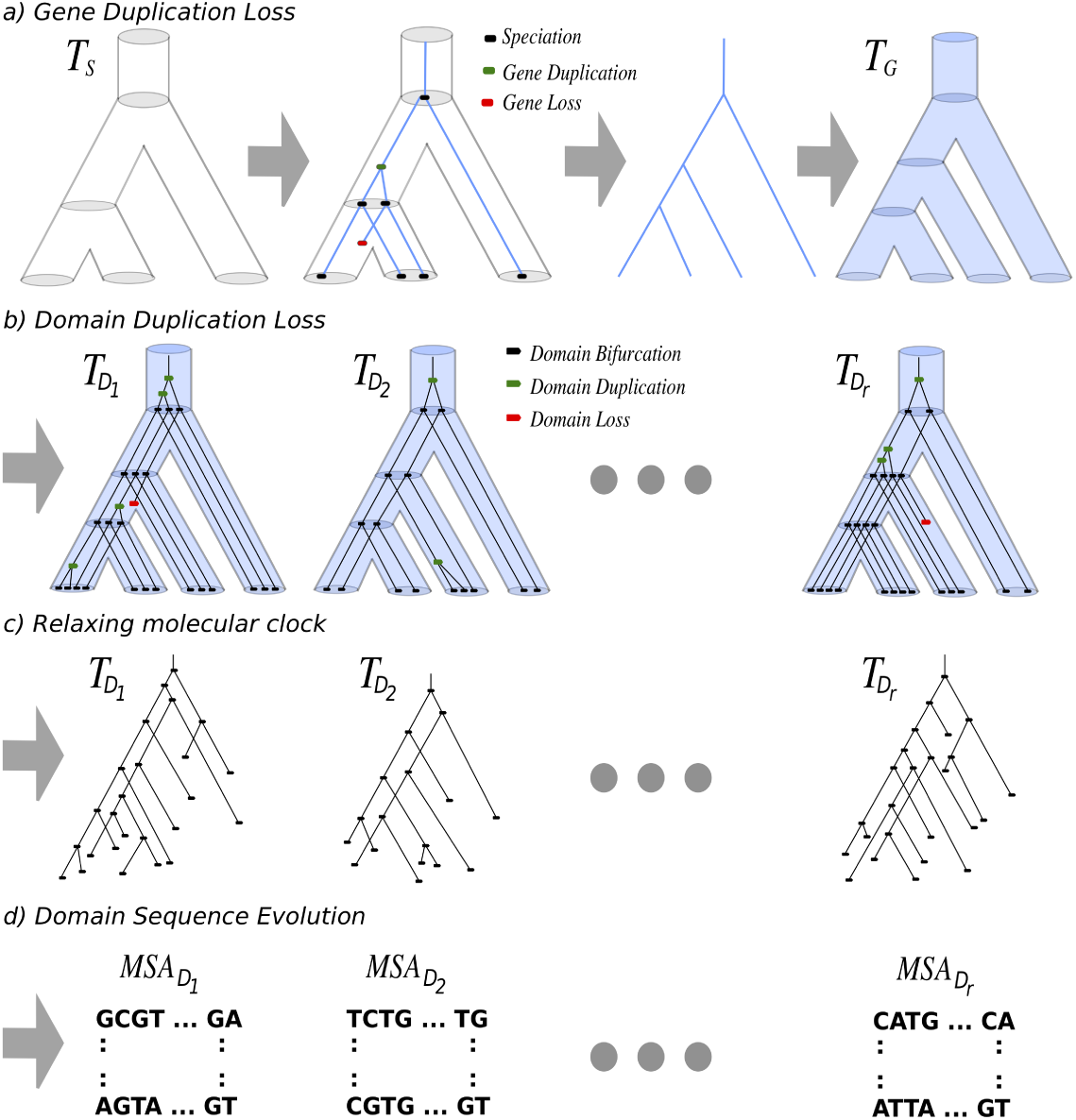
Illustration of the DomainDLRS Model and its submodels. a) A dated species tree *T*_*S*_, *t*_*S*_ is given and a gene tree *T*_*G*_, in blue color, is generated according to the Duplication-Loss model. b) From the dated gene tree *T*_*G*_, *t*_*G*_, a pre-specified number of dated domain trees 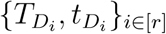 are generated, also using the Duplication-Loss model. c) The molecular clock is relaxed using a rate model, yielding lengths 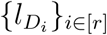. d) Finally, for each domain tree and its lengths 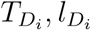, domain sequences are generated for the leaves of 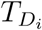 according to a model of sequence evolution.

The Duplication-Loss (DL) model [20] is a birth-death model generalized to occur in a dated tree and describes how a dated guest tree *T*_𝒢_, *t* _𝒢_ evolves inside a dated host tree *T*_ℋ_, *t* _ℋ_. A linear birth-death process with birth rate *δ* and death rate *μ* is used to model duplications and losses over any edge *e* of *T*_ℋ_. The process starts with a single gene lineage at the edge predating the root of *T*_ℋ_ and evolves towards the leaves of *T*_ℋ_. Each time a gene lineage reaches a host tree vertex *x*, it splits into two independent lineages; each evolving down a distinct outgoing edge of *x*. This process continues recursively until it reaches the leaves of *T*_ℋ_. Any vertex of *T*_*𝒢*_ lacking descendants that reached a leaf of *T*_ℋ_ is pruned and any binary vertex that this creates is suppressed, i.e., it is removed and its two former neighbours are made adjacent. In summary, the process results in (i) a binary guest tree *T*_*𝒢*_, (ii) a mapping of the leaves of the guest tree *T*_*𝒢*_ to the leaves of the host tree *T*_ℋ_, and, since the events of the process generating this tree actually takes place in time, (iii) a dating *t*_*𝒢*_ of the guest tree that specifies when a vertex was generated, which together with (i) and (ii) also imply on which host tree edge this happened.

The Rate (R) model [1] is parameterized by a mean *m* and variance *v* and transform the dated, and hence ultrametric, domain trees into length-equipped trees, which typically are non-ultrametric. This is done by, for each edge, perturbing the time associated with the edge to obtain an edge length, which corresponds to the expected number of point mutations over the edge. More specifically, for each edge, an edge length is obtained by taking the product of the edge time and an edge specific rate sampled *i.i.d.* from a gamma distribution with mean *m* and variance *v*.

The formal description of the full DomainDLRS model, generating the dated gene tree, the dated domain trees, and the domain sequences, is as follows: First, a dated gene tree *T*_*G*_,*t*_*G*_ is generated according to the DL model parameterized by the dated host tree *T*_*S*_, *t*_*S*_, duplication rate *δ*_*G*_, and loss rate μ_*G*_. Second, for each *i* ∈ [*r*], a dated domain tree 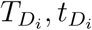, is generated according to the DL model parameterized by the dated host tree *T*_*G*_, *t*_*G*_, duplication rate 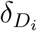, and loss rate 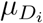. Third, for each *i* ∈ [*r*], an edge length 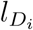 function generated according the R model parameterized by 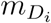 and 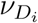. Finally, for each *i* ∈ [*r*], a multiple sequence alignment and a mapping from its rows to the leaves of 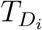 is generated according to M parametrized by 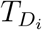 and other possible parameters of *M.*

The DomainDLRS model is, consequently, parameterized by a dated species tree *T*_*S*_, *t*_*S*_, a model of sequence evolution *M*, and

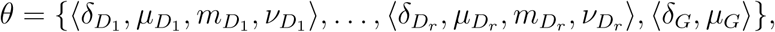

where, for 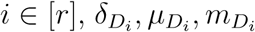, and 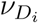, is the duplication rate, loss rate, R model mean, and the R model variation, respectively, for domain *i*, and *δ*_*G*_, *μ*_*G*_ the gene duplication and loss rate, respectively. This is the parameterization used in our analyses. However, we note that, in many cases, it is reasonable, considering to the limited data size, to base an inference on the uniform-domain-rate DomainDLRS model, in which all the domain families have the same rates for the Duplication-Loss as well as the Rate model, i.e., for some *δ*_*D*_, *μ*_*D*_, *m*_*D*_, and *v*_*D*_, for each *i* ∈ [*r*],

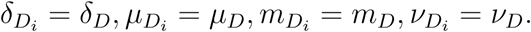

It would be possible to include the domain order in the model and even operations, such as inversions, that rearrange this order. However, focusing on only domain duplications, domain losses, and sequence evolution, allows us to evaluate our result based on how well the result fits the extant domain order. However, when generating synthetic data, we do consider two models that also capture how the domain order evolve over the gene tree: (1) the random model that insert a newly created domain duplicate into a random position and (2) the tandem model that always inserts a new domain next to the original copy.

## 3 Overview of the DomainDLRS-PrIME framework

The three layers of trees in our probabilistic model, i.e., species, gene, and domain trees, render it more complex than earlier models, such as [1, 2, 3]. In order to address the increased complexity, we follow [2] and restrict our attention to most parsimonious reconciliations, thereby effectively giving a probabilistic interpretation to those. Moreover, we introduce a novel technique to sample datings of the vertices that MPR place on edges of a host tree, i.e., either the species tree or the gene tree. This technique allows us to first sample a dating of the gene tree using the dated species tree as host tree and, then, sample a dating for each of the domain trees, now using the dated gene tree as host tree. In contrast to [2], we do not only perform hill climbing over the trees, but investigate three different search strategies: (1) a Grouped Independent Metropolis-Hasting (GIMH) approach [21, 22], where the likelihood of a combination of a gene tree and domain trees is estimated by the average likelihood across a number of sampled datings, (2) a heuristic version of the GIMH, hGIMH, where the likelihood of sampled datings are improved using hill climbing, and (3) a pure hill climbing approach, where hill climbing is performed over all parameters of the model, with the objective of finding a maximum likelihood solution.

The GIMH is a MCMC method that was introduced in [21] and later further explained, as well as generalized to the pseudo-marginal approach [22]. The hallmark of GIMH is that the likelihood of a state is estimated by sampling and the estimate is not updated as long as the state remains the current state. Surprisingly, in contrast to the strategy where likelihood of the current state is reestimated whenever a comparison with the likelihood of a new proposed state is made, the GIMH is guaranteed to converge to the stationary distribution targeted by the Metropolis-Hasting’s ratio used. Although our heuristic version of the GIMH, hGIMH, is not guaranteed to converge to the targeted distribution, it performs superior on synthetic data when evaluated based on how often the estimated MAP tree equals the true tree. It is also superior to the pure hill climbing, which actually is better that the GIMH, see figure 2, below. For this reason, the hGIMH sampler is selected to be our DomainDLRS method.

The input of the hGIMH sampler, as well as the two other methods, is a dated species tre*e* 〈*T*_*S*_,*t*_*s*_〉, where *T*_*s*_ is the actual tree and *t*_*s*_ the dating of its vertices, and a multiple sequence alignment for each domain family, i.e., different multiple sequence alignments for different domain families and with a number of rows that may vary across domain families. It also requires a labelling of each gene with the species in which it is found as well as a labelling of each domain with the gene in which it is found. Let *T*_*G*_ denote a gene tree and let the *i*:th domain tree be denoted by 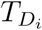 and its edge lengths by 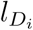. Let *θ* denote additional parameters of the model, such as the parameters of *M* and the rate parameters. A state in our sampler is a tuple

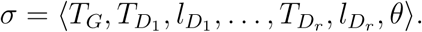

When in current state *σ* and the new state σ ^’^ is proposed, the sampler’s MH ratio is given by

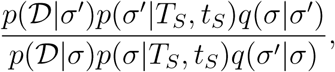

where *q*(·*|*·) is the proposal distribution. The proposal distribution, which is mostly composed of standard proposal distributions, as well as how to compute proposal probabilities *q*(*·*·*|·*·) are described in Supplementary material. Moreover, the probability of the data *p*(*𝒟 | σ*) can easily be computed using Felsenstein’s so-called pruning algorithm [23]. So, the main technical difficulty involved in computing the MH ratio consist of evaluating *p*(*σ |T*_*S*_, *t*_*S*_): the probability of the gene tree and the length equipped domain trees, given the dated species tree; a task performed by sampling datings and applying a hill climbing approach.

Our strategy builds on a two-layered sampling approach to estimate *p*(*σ|T*_*S*_, *t*_*S*_). First, notice that a dating *t*_*G*_ is either incompatible with the dating *t*_*S*_ or the datings imply a reconcilation of *T*_*G*_ and *T*_*S*_. Let *M* be of set of datings *t*_*G*_ that together with *t*_*S*_ imply the MPR of *T*_*G*_ and *T*_*S*_. A dating *t*_*G*_ is sampled from *M* according to the unique distribution over *M* that is proportional to *p*(*t*_*G*_*|T*_*G*_,*T*_*S*_, *t*_*S*_, *σ*). Second, for each domain tree 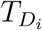, a dating 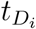 is sampled analogously, that is, (1) let *M*_*i*_ be of set of datings 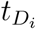 that together with *t*_*G*_ imply the MPR of 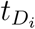 and *T*_*G*_ and (2) sample a dating 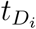 from *M*_*i*_ according to the unique distribution over *M*_*i*_ that is proportional to 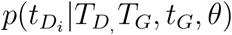. The datings, *t*_*G*_ and 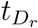 for each domain family *r*, obtained from these two steps yields the estimate

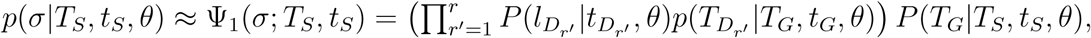

where *P*(*T*_*G*_*|T*_*S*_, *t*_*s*_, *θ*) and 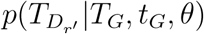 are the probabilities of *T*_*G*_ and 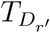, respectively, according to the DL process and 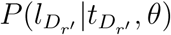 is the probability of the rates induces by 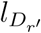, and 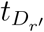, To obtain improved estimates of *p*(*σ |T*_*S*_, *t*_*s*_, *θ*), we use averaging over multiple samples as well as hill-climbing techniques. The details of the the two sampling steps are described in Supplementary information Section **??** andSection **??**.

## 4 Results

In this section, the performance of DomainDLRS on synthetic as well as biological data is evaluated as well as compared with that of a more conventional two-step approach, MrBayesMPR, described below.

### 4.1 MrBayes-MPR

We will below compare the performance of DomainDLRS with a method we call MrBayesMPR, which formalize, what could be called, the ‘standard’ approach of independent reconstruction of domain trees, followed by computation of MPR between domain tree and gene tree. We use MrBayes to estimate the posterior distribution of domain trees and gene trees. The domain trees are estimated from aligned sequence data from the individual domains and gene trees are estimated from the full gene alignments. Consequently, it is reasonable to expect that, for MrBayes, the accuracy achieved when reconstructing gene trees gene from data generated under the tandem model should be better than that achieved when data is generated from the random model. In contrast, DomainDLRS is oblivious to the choice between the random and the tandem model, since it relies on aligned domains and is insensitive to the order in which those appear in a gene. For MrBayes-MPR, we use MPR to reconcile the MAP domain tree with the gene tree.

### 4.2 Synthetic data analysis

The accuracies of our methods as well as that of MrBayes-MPR were evaluated on semisynthetic data, derived from three gene families ZNF91, ZNF468, and ZNF679. We first estimated the posterior distribution over the parameters of the DomainDLRS model using hGIMH and then, based on the MAP parameters obtained, generated three sets of synthetic gene families here referred to as SZNF91, SZNF468, and SZNF679, each containing 100 synthetic gene families. The accuracy of each inference method was then evaluated based on how well the true synthetic domain trees and the true synthetic gene trees were reconstructed. For domain trees, the evaluation was concerned with both the accuracy of domain duplication identification and Robinson-Fould (RF) distance between the reconstructed domain tree and the true domain tree.

**Figure 2:**
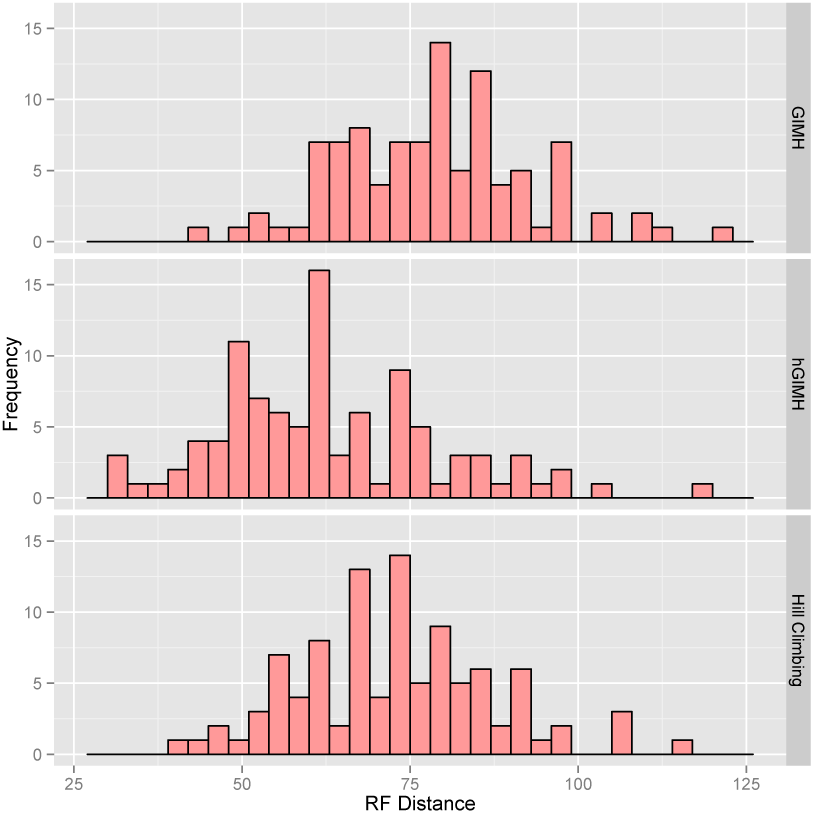
The Robinson Foulds (RF) distance distributions on synthetic data for our three candidate methods. The first panel, from the top, shows the RF distance distribution of GIMH, the second panel shows RF distance distribution of hGIMH, and the third panel shows the RF distance distribution of the Hill Climbing method. It is clear that hGIMH performs better than the other two methods.

We first evaluated our three DomainDLRS methods, Hill Climbing, hGIMH, and GIMH, based on how well they reconstructed the true synthetic trees in the SZNF91 data set. The result, shown in Figure 2, shows that the hGIMH method performed clearly best, dispaying the lowest RF distance distribution. Of the remaining two methods, Hill climbing performed slightly better than GIMH. For this reason, hGIMH is the method that below will be compared with DLRS and MrBayes and we will, henceforth, refer to it simply as DomainDLRS.

**Figure 3:**
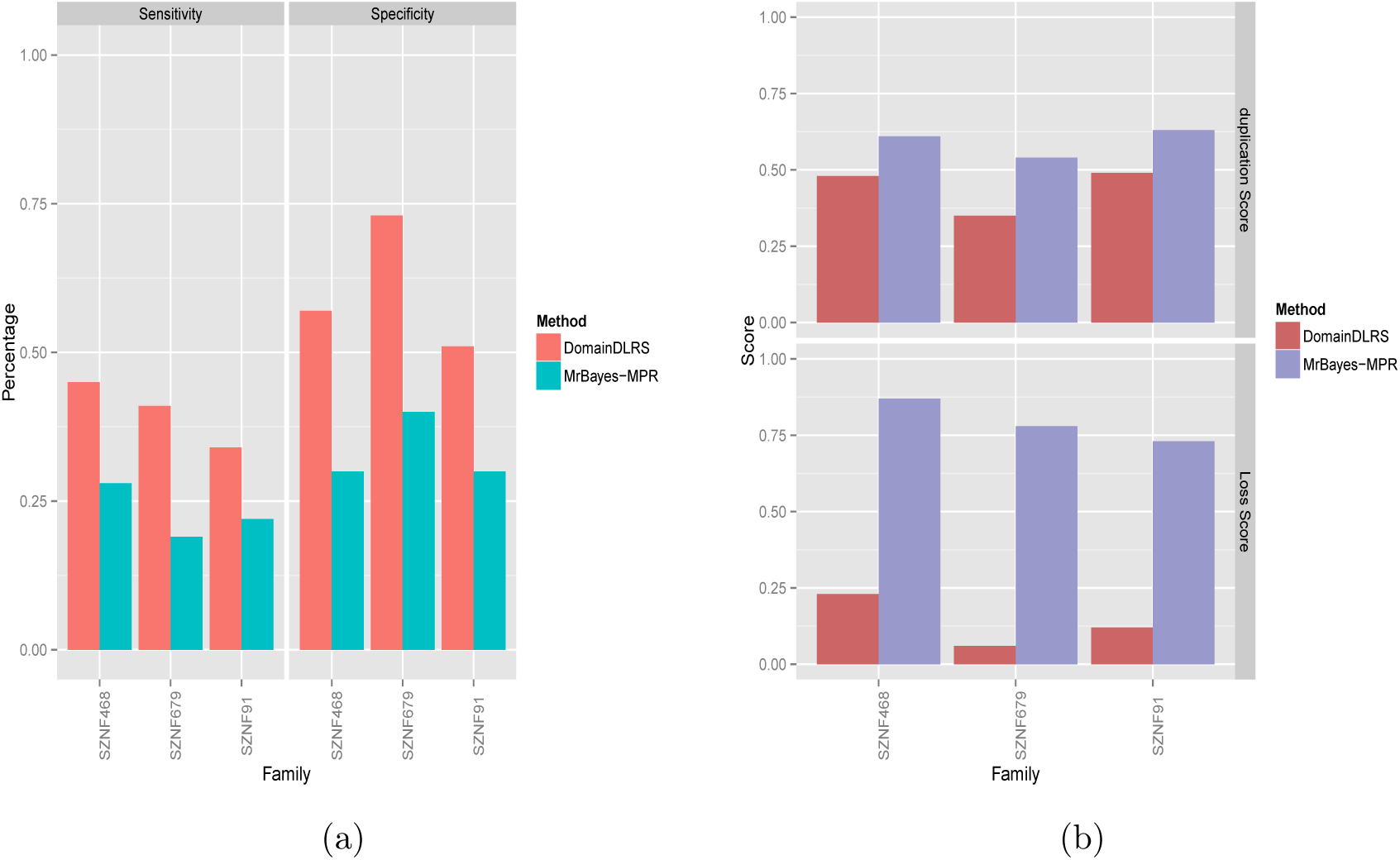
Accuracy and duplication-loss scores of DomainDLRS and MrBayes-MPR over synthetic datasets of SZNF91, SZNF468, SZNF679. a) The bar chart shows the sensitivity and specificity of domain duplication events labelled by DomainDLRS and MrBayes-MPR methods for synthetic datasets. b) The second bar chart shows the duplication and loss scores of DomainDLRS and MrBayesMPR.

Secondly, we compared DomainDLRS to MrBayes-MPR to evaulate their relative performance. For the three synthetic data sets, both the specificity and sensitivity are clearly better for DomainDLRS than for MrBayesMPR, approximately improving both statistics substantially, in several cases by a factor of approximately two, Figure 3(a). DomainDLRS also performs substantially better when comparing the RF distances, Figure 4. DomainDLRS clearly performs superior to MrBayesMPR. It is also interesting to note that MrBayes has an alarmingly high loss score, Figure 3(b)

**Figure 4:**
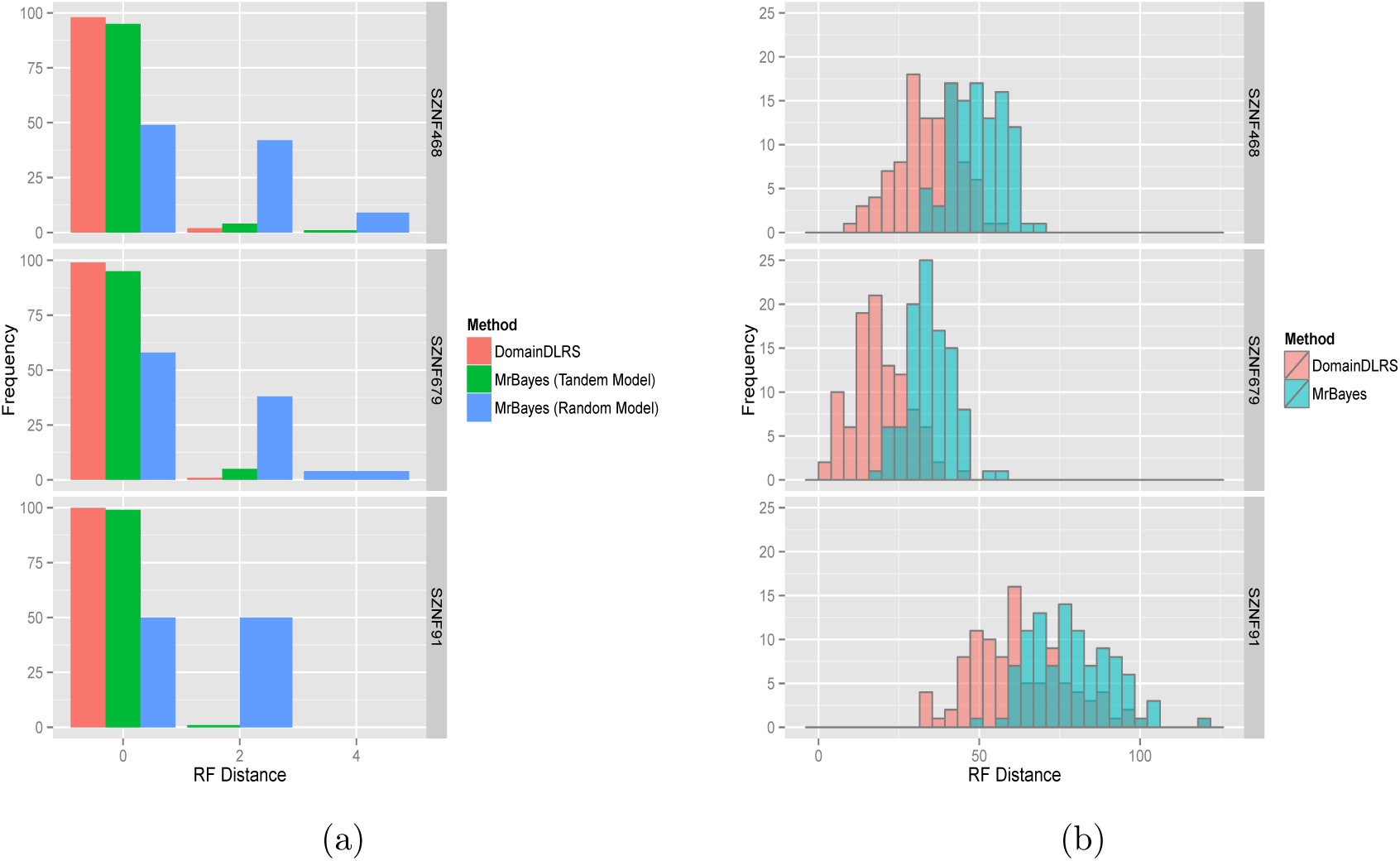
Distribution of Robinson Foulds (RF) distances of inferred gene and domain trees over synthetic datasets of SZNF91, SZNF468, SZNF679. a) A comparison of RF distance distribution of gene tree inferred by DomainDLRS and MrBayes, respectively, to the true gene trees. For MrBayes, we distinguish between data generated under the random and the tandem model to capture the expected performance difference. b) The bar chart shows the comparison of RF distance distribution of domain tree inferred by DomainDLRS and MrBayes based on the synthetic true domain trees for SZNF91, SZNF468, SZNF679 datasets.

We then evaluated the ability to reconstruct the *gene tree* – which is considerably smaller than the domain trees, so the RF distance is most often 0 or 2. To construct gene input data for MrBayes, we concatenated the generated domain sequences according to a domain order. This domain order was generated according to two different models; for each new copy generated by a domain duplication, the *random model* selected a random insertion point in the currently existing domain order, while the *tandem model* inserted it next the original domain. Notice that DomainDLRS takes individual domain sequences as input and ignores domain order. It turns out that MrBayes performs very poorly for data generated according to the random domain order model (Fig. 4). In contrast, the performance of MrBayes is very good for data generated according to the tandem domain order model, in fact, almost as good as for DomainDLRS.

### 4.3 Biological data analysis

We applied our framework to the Zinc Finger gene families ZNF91, ZNF468, ZNF558, ZNF611, ZNF679, and ZNF764 in four primate species, *Macaca mulatta, Pongo abelii, Pan troglodytes* and *Homo sapiens.* These families are both methodologically challenging as well as biologically interesting because, in each family, the number of Zinc Finger domains varies across its members, a phenomena which which is likely to be caused by domain duplication and loss. Some of these families have also been exposed to recent primate-specific gene duplications. We present the results from the comparison between DomainDLRS and MrBayesMPR, first on gene tree level, before focusing on domain tree level.

#### 4.3.1 Gene evolution

For four of the gene families (ZNF468, ZNF679, ZNF764/747 and ZNF91) DomainDLRS and MrBayesMPR reconstruct the same gene tree, while for the remaining two (ZNF611/600 and ZNF558/557) the two methods disagree. For ZNF611/600, DomainDLRS indicates that the Macaque ZNF611 gene is not the sister to the other primate ZNF611, but is instead the product of an ancient gene duplication (Supplementary Fig. **??**). This is consistent with the well supported domain tree, where a number of shared domain duplications supports the closer relationship between the ZNF611 and the hominid ZNF600 subtrees (Supplementary Fig. **??**). This shows how taking individual domain evolution into account can improve our understanding of gene family evolution. Also for the ZNF558/557 gene family, the gene tree estimate from MrBayesMPR differs from that from DomainDLRS (Supplementary Fig. **??**), and again the latter is corroborated by a well-supported domain tree (Supplementary Fig. **??**). Here, the MrBayesMPR gene tree is probably affected by a long branch attraction between Macaque ZNF557 and Pongo ZNF558 genes together with erroneous rooting of the gene tree. Notice that MrBayes reconstructs an unrooted gene tree, which need to be rooted by some external criterion. Here, this rooting is performed using Notung, which attempts to find the rooting that minimizes the number of duplications and losses. However, branch length information from the sequence data is not used, adding another level of ignored uncertainty information to the final MPR. In contrast, rooting of the gene tree is an integrated part of DomainDLRS and different rootings are evaluated by their probablility under the DomainDLRS model, hence, making use of all available information. As the gene trees reconstructed by DomainDLRS appears to be more correct we will, to enhance comparison of domain evolution, use the Domain DLRS domain trees as the (host) gene tree for the MrBayesMPR method when analyzing domain evolution.

#### 4.3.2 KRAB domain evolution

In almost all gene families, the evolution of the KRAB domain, follows the gene tree exactly. The exception is ZNF611/600, in which the KRAB domain is lost from two of the gene paralogs in all species. However, these two losses correspond to two separate loss events in a domain tree that otherwise is congruent with the gene tree.

**Figure 5:**
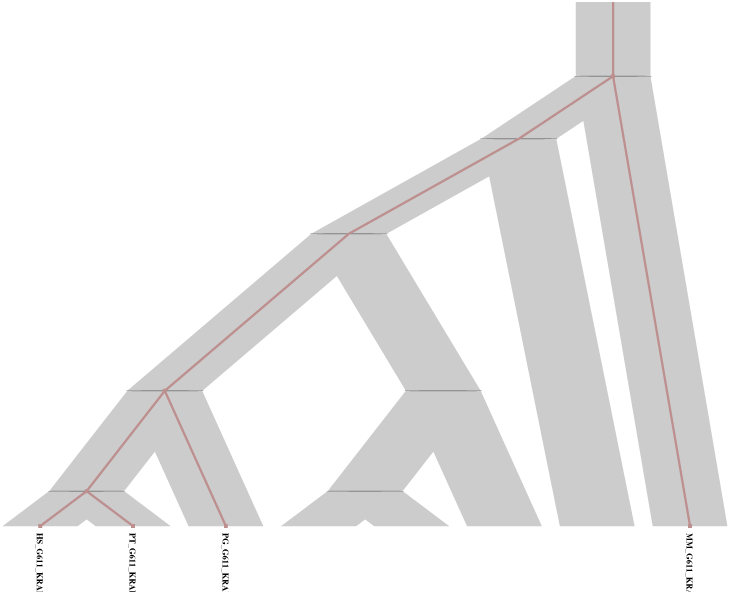
The reconciliation of KRAB domain tree of ZNF611 family with the gene tree inferred by DomainDLRS.

#### 4.3.3 ZNF domain evolution

##### Recent and ancient duplications

Our framework provides datings of the domain tree vertices and we will, for enhancement of results description, distinguish between recent and ancient domain duplications (and consequently also between recent and ancient domain tree vertices/edges), where we define *ancient* as older than the root vertex of the gene tree. We investigated the posterior probability distributions for ancient and recent domain tree edges in our ZNF domain families (Fig. 6). Interestingly, in the DomainDLRS and in particular the MrBayesMPR results, there is a substantial enrichment for low posterior probabilities among ancient edges. In fact (1) many ancient edges have low posterior probabilities, on average 34% of the ancient vertices in the DomainDLRS results have a posterior probability < 0.8, but the corresponding figure for mRBayesMPR is as high as 56% and (2) for DomainDLRS on average 1.6% recent edges have posterior probability < 0.8, but the corresponding figures for mRBayesMPR is as high as 21%

**Figure 6:**
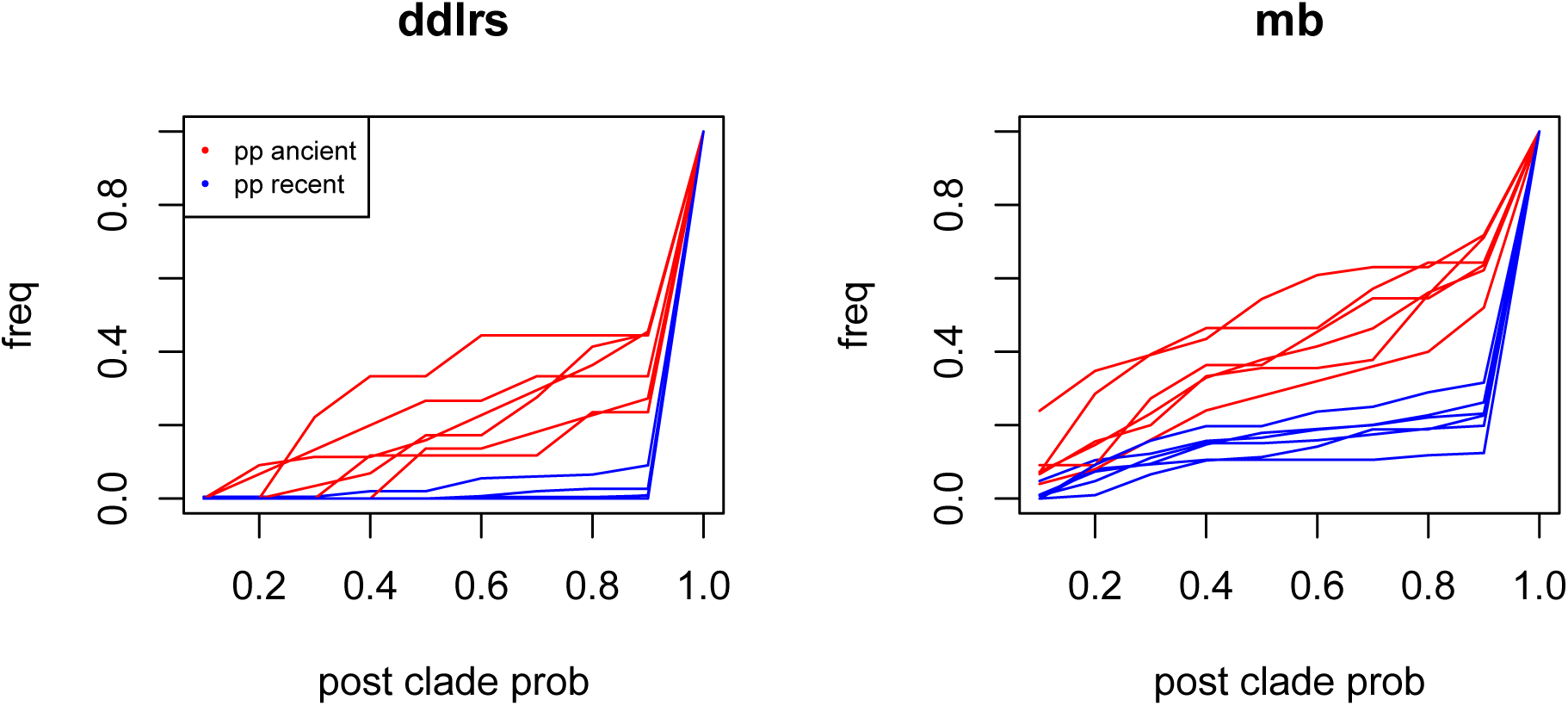
Clade posterior probability cumulative frequency distribution of recent(red) and ancient (blue) vertices for (a) DomainDLRS and (b) MrBayes-MPR over all biological datasets.

A possible explanation for this are the constraints implied by the gene tree for events occurring above and below its root, respectively. For recent domain vertices, below the root, the gene tree topology provides rather fine-grained constraints on the timing of domain events, which therefore can be robustly estimated. In contrast, the single edge leading to the root provides much less time constraints and the reconstruction of the time of ancient domain tree vertices (which may be considerably older than than the root) is inheritantly much harder, in practice requiring additional sampling of taxa/genes to break up the root edge. Because of this, we will mostly focus on recent domain tree edges in our discussion of the present results.

**Figure 7:**
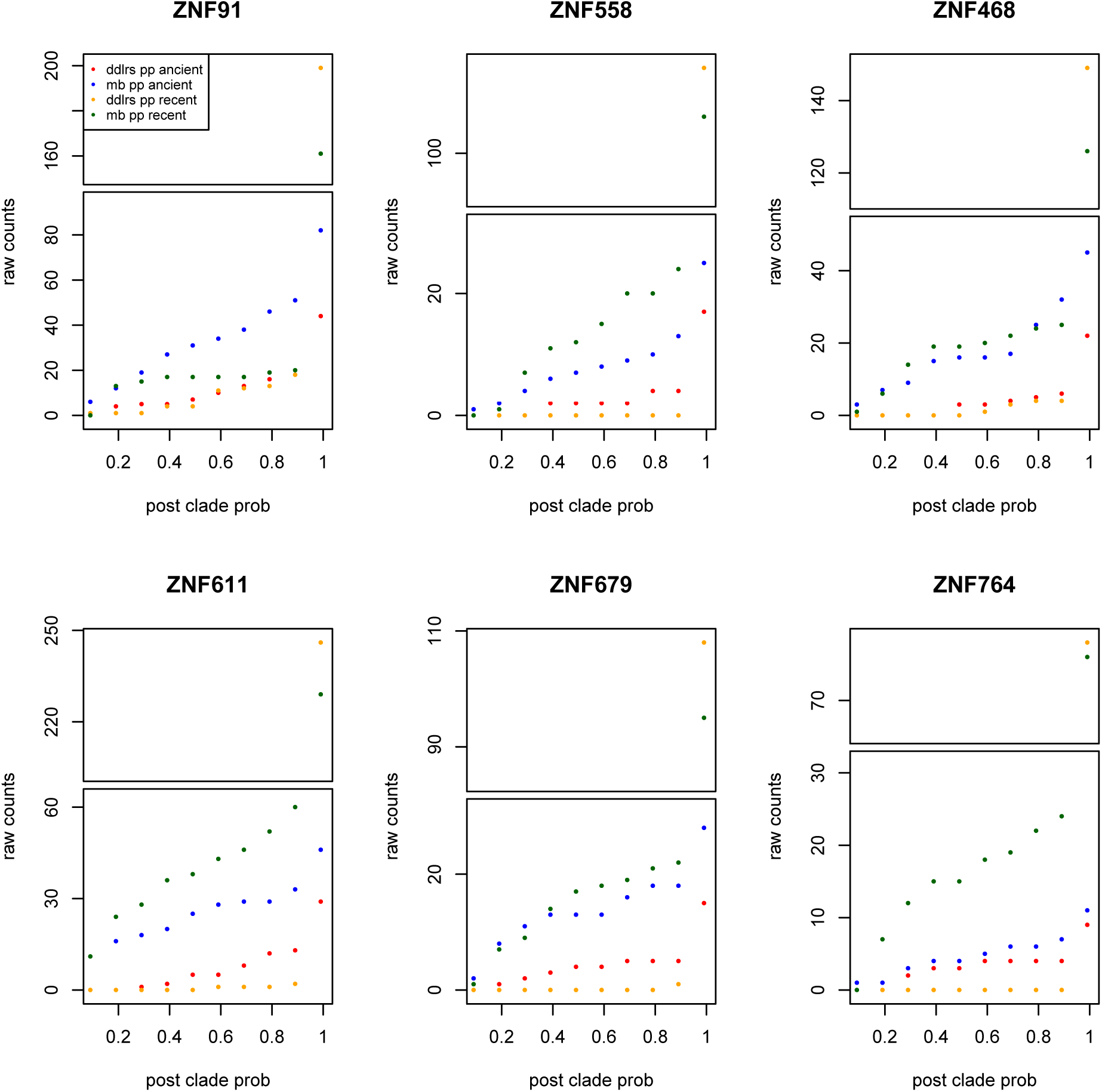
Clade posterior probability cumulative raw counts distribution of DomainDLRS (red) and MrBayes (blue) for each biological datasets.

#### 4.3.4 Robustness

For all ZNF-domain families, it is clear that the DomainDLRS results provides a clearer and more well-supported picture of the domain evolution than do MrBayesMPR. Firstly, MrBayesMPRDomainDLRS has considerable higher support for all vertices, and in particular for recent vertices (Fig. 7). While the cumulative count distribution of DomainDLRS posterior clade probabilities, for almost all domain families, is strongly peaked at posterior probability 1.0, with only very few or no vertices at lower posterior probabilities (this is most pronounced for recent vertices), MrBayesMPR displays more flattened distributions with a large fraction of vertices at lower posterior probabilities. The effect can clearly be seen in the reconciled MAP trees for ZNF764 (Fig. 8), which have relatively few domains, as well as few domain events, where DomainDLRS gives an extremely clear view of the very well supported domain tree, while MrBayesMPR produces a much more weakly supported domain tree that reconciles badly with the gene tree (Fig. 7). However, it is also evident, also in more complex reconciled MAP trees (Supplementary Figs. **????**), for which DomainDLRS consistently provides better supported domain tree vertics (Fig. 7 and Supplementary Tables **??**-**??**).

**Figure 8:**
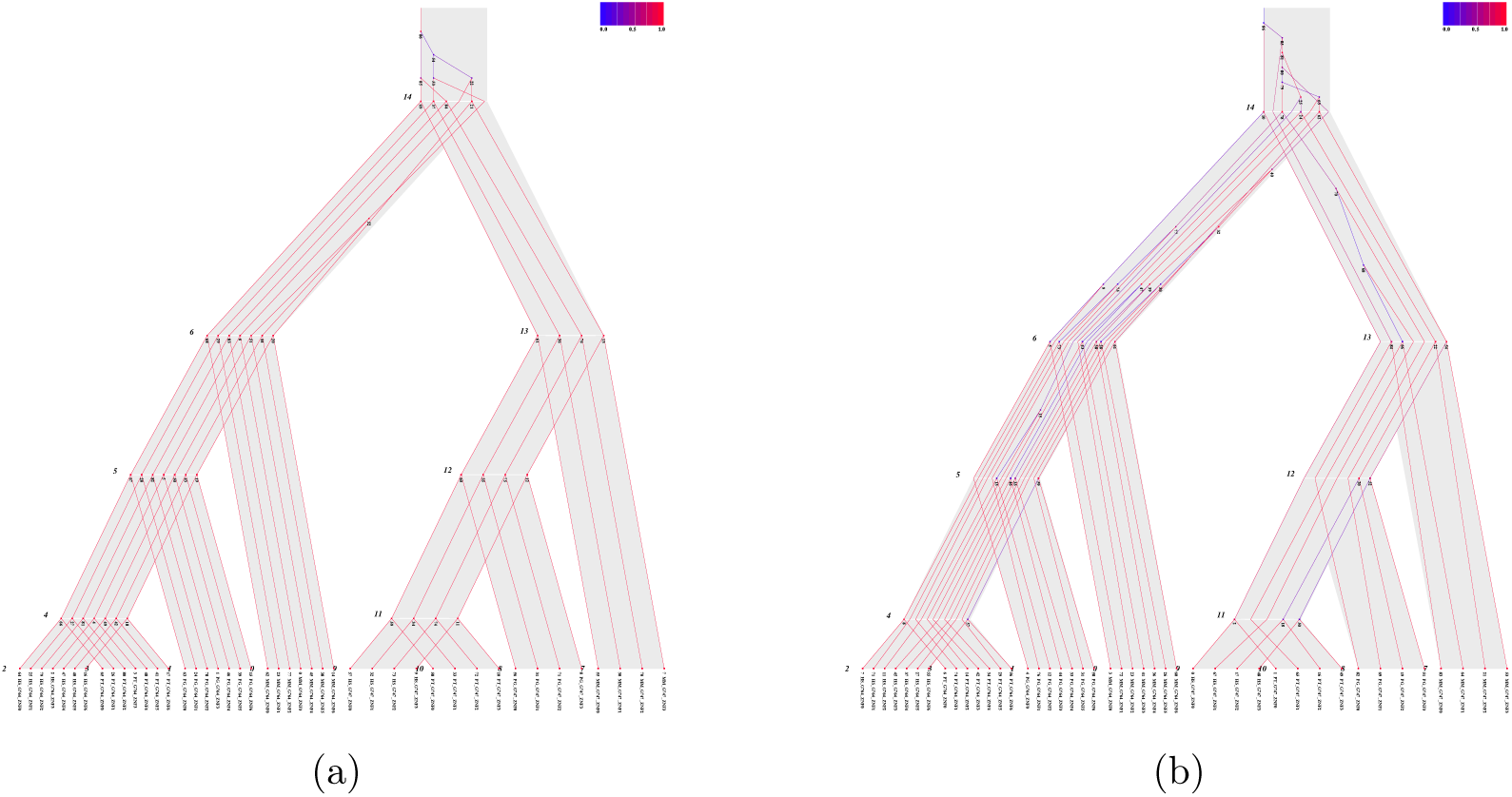
The most parsimonious reconciliation of domain tree with gene tree for gene families ZNF764 inferred by DomainDLRS (a) and MrBayes (b).

Secondly, DomainDLRS provides much more clear and convincing picture of domain evolution and particularly of recent domain evolution (Supplementary Figs. **??**-**??**).

In general, DomainDLRS manage to explain more domain vertices as implied by, and coinciding with, gene tree vertices (henceforth referred to as *domain bifurcations*), and thereby requiring substantially less number of domain duplications and domain losses compared to MrBayes-MPR (Fig. 9).

**Figure 9:**
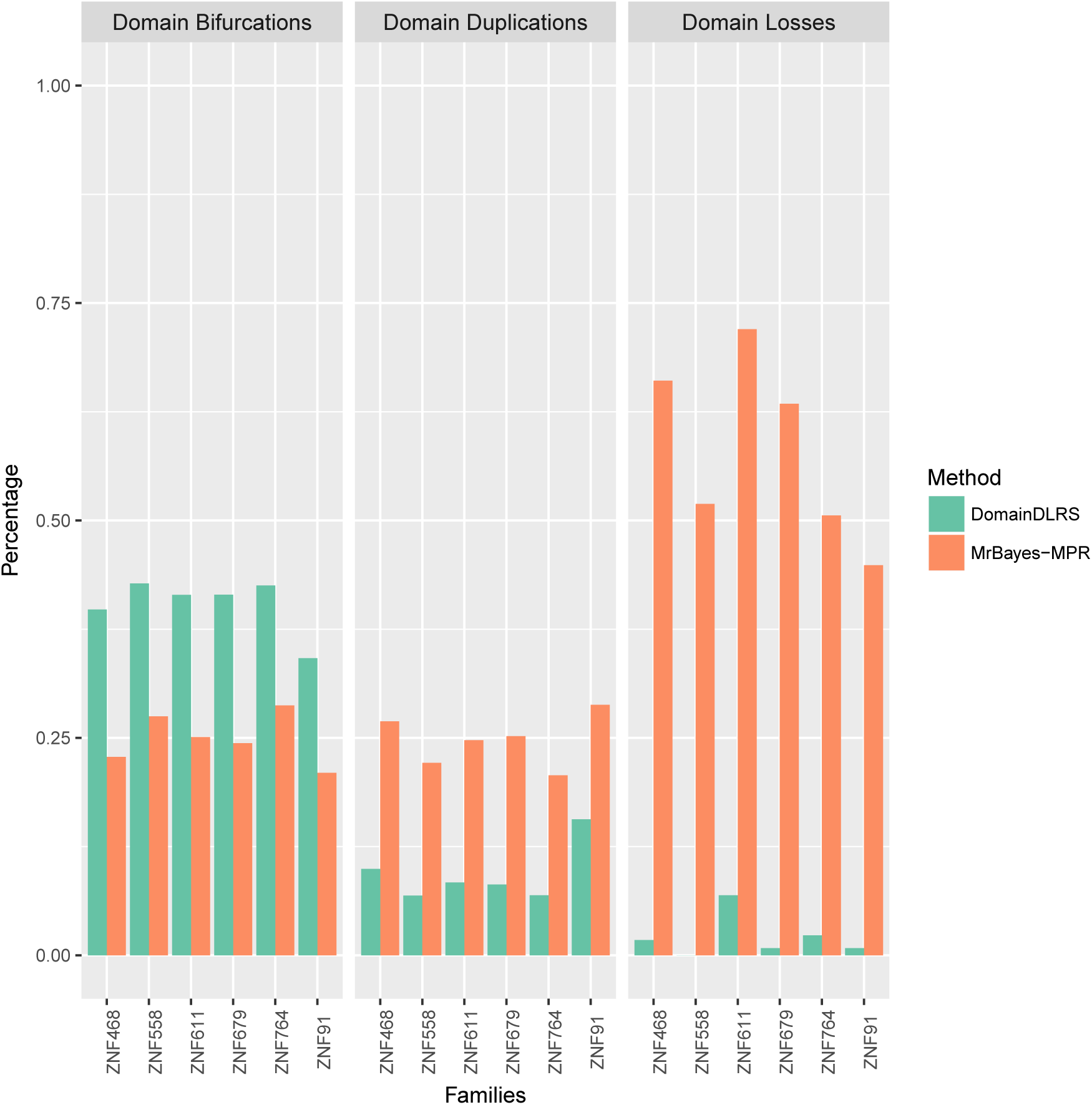
The figure is showing percentages of domain duplications, domain losses, and domain bifurcations implied by gene duplication and speciation events with respect to total number of vertices of the domain tree in all mentioned biological datasets.

**Figure 10:**
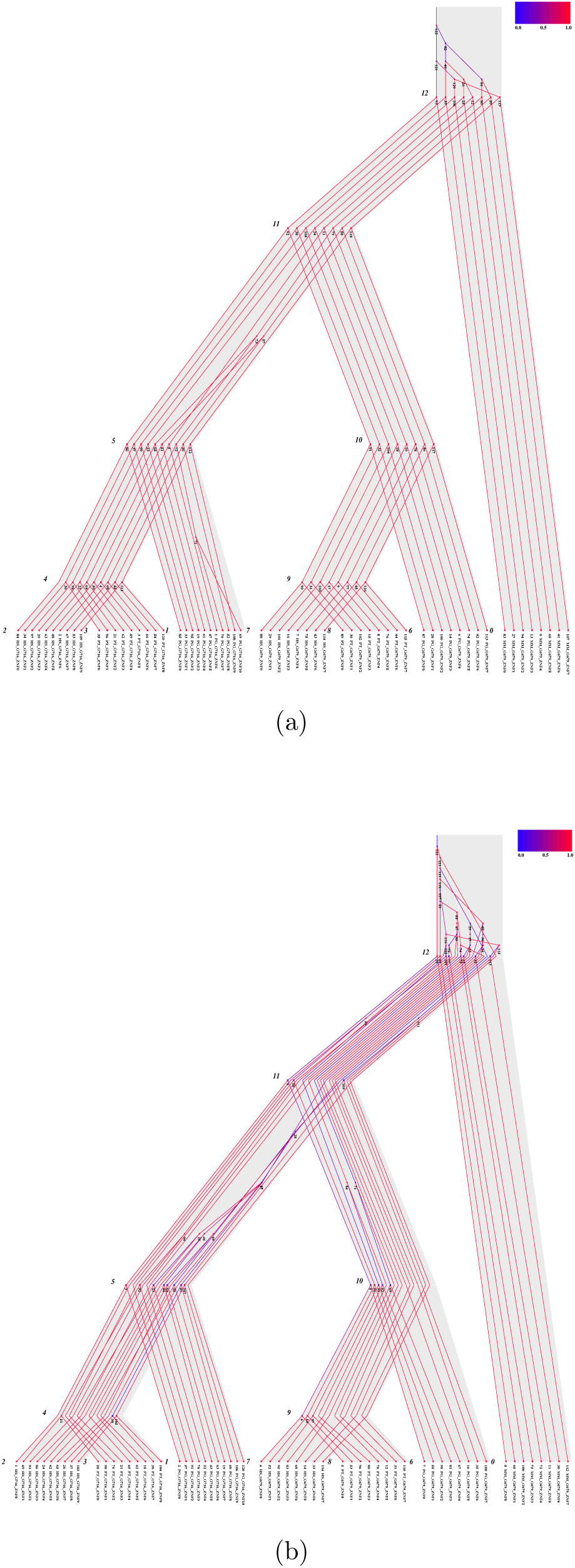
Reconciliation heat maps of ZNF679 Zinc-Finger domain trees inferred by a) DomainDLRS and b) MrBayes-MPR. It shows a reconciliation between a domain and a gene tree with domain tree leaves ordered as the domains occur in the genes and each domain tree edge coloured according to its posterior probability.

Thirdly, for all domain families, MrBayes-MPR predicts more nodes as ancient compared to DomainDLRS (Supplementary Table **??**). This is a well-known indication of a reconciliation of an incorrect tree (in this case the domain tree) [24]; given the enrichment of low clade support for MrBayes, the MrBayes domain tree is very likely to contain weakly supported topology incongruences with the gene tree, such that MPR forces domain vertices to be placed higher up in the tree (Fig. 10(b)). Because the MrBayes domain tree reconstruction and MPR are performed as two separate steps, the MrBayes posterior clade probabilities are not considered in the MPR, resulting in ‘false’ ancient nodes. DomainDLRS, on the other hand, which integrates tree reconstruction and reconciliation, balances probabilities obtained from sequence evolution and DL models and provides a more robust result. Moreover, as discussed above for gene tree recnstruction, in DomainDLRS the rooting of the domain tree is integrated into the analysis, while MrBayesMPR, by design, must rely on a some external rooting method. This is obviously an advantage of DomainDLRS, as an erroneous rooting of the domain tree will conflate the problem of surplus ancient vertices.

Lastly, DomainDLRS provides ancestral genes with more realistic domain contents (Supplementary Tables **??**-**??**). For example, in MrBayes reconciliation of ZNF91, the gene in the least common ancestor (LCA) of human and Old-World monkeys had 59 Zinc-Finger domains (Supplementary Table **??**). This is far more than the 34 domains that is the maximum number of Zinc-Finger domains in any of the extant genes in the family (implying that 109 domain losses have occurred, Supplementary Table **??**) For DomainDLRS, the number of Zinc-Finger domains in the LCA is 23 and the number of losses 0 (Supplementary Table **??**). This pattern is consistent for all ZNF gene families.

#### Domain order

Notice that when reconstructing domain trees, DomainDLRS (or MrBayesMPR) does not take advantage of domain order information. However, domain order information within the gene can be used to evaluate the realism of the reconstructed domain trees. We have used a parsimony-based algorithm to draw the reconciled trees. The algoritm greedily orders domain lineages on internal vertices with the objective of minimizing the number of crossing domain tree edges given the known domain orders in extant genes (Figs. 8, 10 and Supplementary Figs. **??**). In the reconciled trees, domain shuffling events (and also duplications) can be seen as crossing-over of lineages. The result is not guaranteed to be perfect, and can often be improved manually (e.g., 11(a)), but allows a rough estimate of domain shuffling events in the genes evolution. Focusing here on the DomainDLRS results, we see that overall few domain shuffling events are predicted; the exception is domain-rich ZNF91 family, where domain shuffling appears to have occurred in the ancestor of Homo and Pan and in the Pong and Macaca lineages. It is clear, for all gene families, that the domain order implied by DomainDLRS reconciliations for inner vertices are more consistent with the domain order in extant genes than those of MrBayesMPR.

#### Segmental duplications

Inspection of the reconciled trees and the inferred gene ordering also allows us to study the relative numbers of single and segmental duplications. Using a rough definition, requiring that segmental duplications (1) occur on the same edge, (2) can occur at the same time (a vertex and its ancestor cannot be in the same segmental duplicaiton), and (3) is consistent with the inferred domain ordering without requiring too many structural mutations, we estimated the number of single and segmental duplications (Table 1). While there are approximately the same number of single and segmental duplications, the number of domains affected by segmental duplications is naturally higher than the number of domains affected by single duplicatins.

**Table 1:**
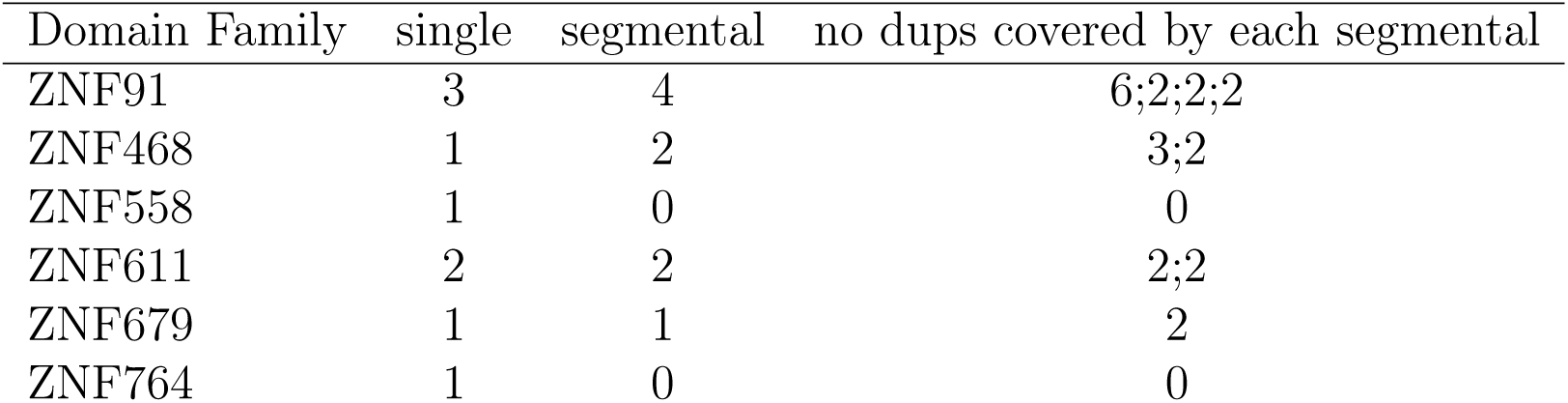
Estimated number of segmental duplications from the DomainDLRS analyses of the biological datasets.

### 4.4 A case study: ZNF91

The ZNF91 family appears to have emerged in the LCA of humans and Old-World monkeys and published phylogenetic analyses indicate that it have undergone dramatic structural changes, including the addition of seven Zinc Fingers domains in the LCA of humans and chimpanzees, through segmental duplication [5]. Moreover, the study of Jacobs et al. [5] also suggested that the ZNF91 evolution is driven by an evolutionary arms race with retrotransposons. Their phylogenetic analysis of the ZNF91 family was based on the assumption that Zinc-Finger domains 7-12 in the human, chimpanzee, and gorilla genes has been created by a segmental duplication of domains 18-23 in the ancestor of these species (the hominine ancestor) causing the human, chimpanzee and gorilla to have substantially more Zinc-finger domain than the other species involved in their analysis. Additionally, they assumed, based on perceived similarity between Zinc-finger 6 and 7 in human, that these two domains are the result of a more recent than their proposed segmental duplication.

However, because DomainDLRS directly models the evolution of the individual zinc finger in ZNF91, it allows show that the assumption underlying their analysis is incorrect as well as provide a more refined analysis, in general. It follows from our DomainDLRS analysis (Fig. 11) that the segmental duplication event occurring in the hominine ancestor actually created Zinc-Finger domain 4 and domains 6-10 from domain 16-21 (while we have not sampled Gorilla here, this event is likely to correspond to the segmental duplication event of Jacobs et al.). This means that the ancestor of human zinc finger domains 6 and 7 predates the root of the hominine tree in our analysis and, thus, there is no support for a recent duplication producing these domains. Moreover, domain 4 appears to be the result of a domain duplication followed by a domain loss. In fact, the ancestral domain order can be read directly from the reconciled domain tree in Figure 11. In summary, DomainDLRS directly provides the ancestral domain content, and combined with an algorithm for inferring domain order at internal vertices, constitutes a powerful tool for evolutionary genomics studies such as the one by Jacobs et al. [5].

**Figure 11:**
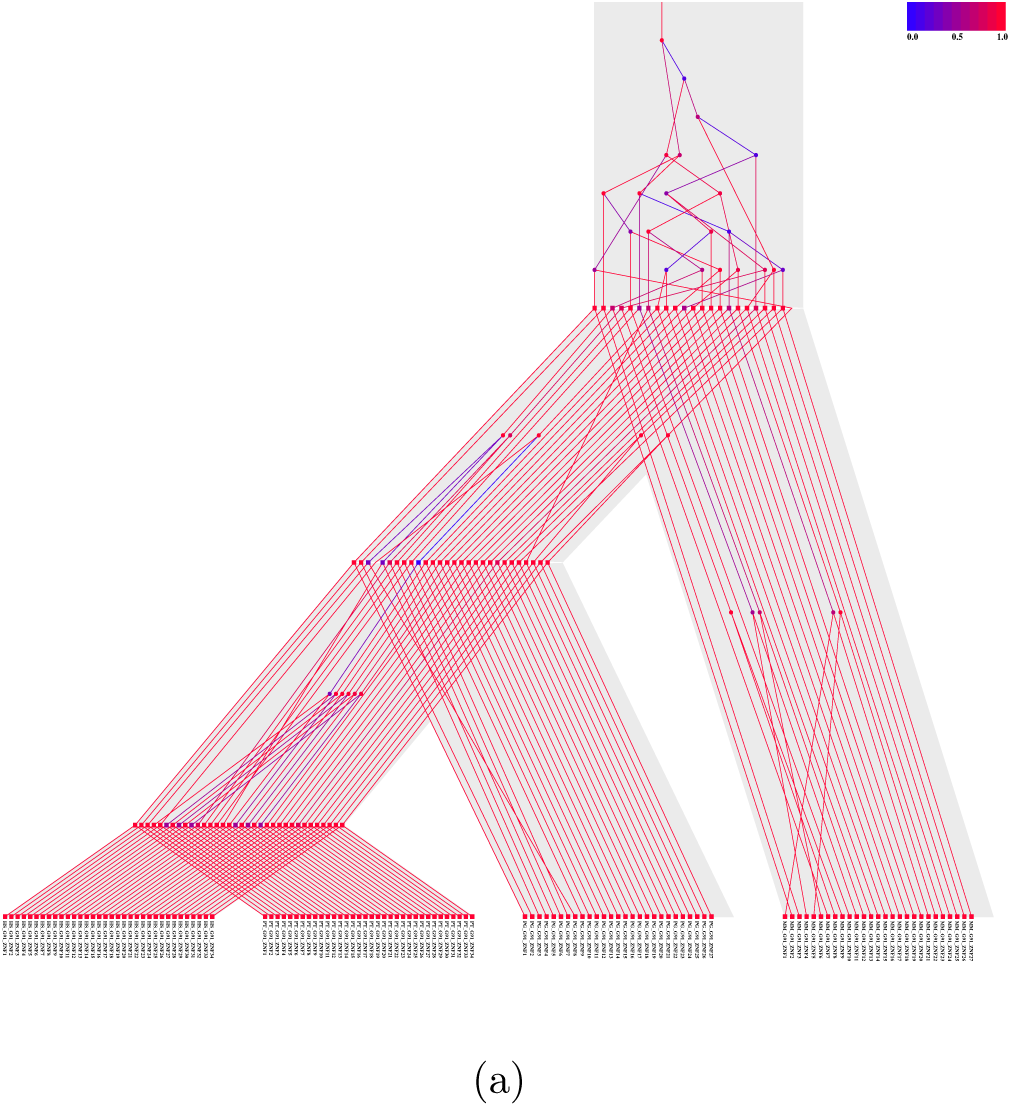
Reconciliation heat maps of ZNF91 Zinc-Finger domain trees inferred by DomainDLRS. It shows a reconciliation between the domain and the gene tree with domain tree leaves ordered as the domains occur in the genes and each domain tree edge coloured according to its posterior probability. Domain ordering on internal nodes have been improved manually.

## 5 Discussion

We have presented a hierarchical model of domain evolution including a nested version of the Duplication-Loss process, with three levels of trees, which describes how a gene tree evolves inside a species tree and how several domain trees evolves inside this gene tree. We have also presented and evaluated the three inference frameworks for this model and concluded that among these DomainDLRS, a heuristic version of a GIMH algorithm, performs best. Our inference framework takes a dated species tree and aligned *domain* families as input and provides a posterior over tuples of each containing a dated gene tree reconciled with the species tree and dated domain trees reconciled with the gene tree.

Our framework provides several advantages compared to the traditional approach, where tree reconstruction and reconciliation are performed independently. Firstly, taking aligned domain sequences rather than gene sequences as input, constitute a tremendous advantage for gene tree reconstruction. From a technical viewpoint, direct alignment of genes that have undergone multiple, and in many cases non-tandem, domain duplications is a hard unsolved computational problem. This problem is here completely evaded. Secondly, simultaneous modeling of tree reconstruction and tree reconciliation balances probabilities obtained from sequence evolution and duplication-loss models and, thereby, gives more stable and accurate pictures of domain (and gene) evolution. Thirdly, our integrated framework also provides rooting of both the gene and domain trees.

We compared DomainDLRS with MrBayesMPR, a method that combines a MrBayes inference of a domain trees and a MPR reconciliation of the domain tree and a gene tree. It is clear, both from our analyses on synthetic data and on biological data from several zinc-finger gene families in primates, that DomainDLRS provides much better gene and domain tree reconstruction than MrBayesMPR.

In the analyses of synthetic data, DomainDLRS clearly performs better than MrBayesMPR both in terms of sensitivity, specificity, and similarity, in terms of RF distance, of the reconstructed trees to the known domain and gene trees. It may at first sight seem possible to argue that, at least for the synthetic SZNF91 families, the RF distance might be too high to render the biological analysis in Section 4.3 entirely trustworthy. The biological analysis, however, stand out as robust in its own right, see Section 4.3. One reason for the relatively lesser performance in the synthetic case may be that for the biological data a fairly high number of DomainDLRS duplications precedes the root and are of less significance, while for the synthetic data the domain duplications appear everywhere and all of them can be considered to be equally important. We also generated synthetic gene data under two different models for domain duplication either with random or with tandem insertion of the new copy in the gene. We show that MrBayesMPR clearly performs worse under the random model than under the tandem model (DomainDLRS is not affected). The analyses of biological data in reveal that the appropriate model lies somewhere in between these two extremes, although closer to the tandem than the random domain order model. It is natural to conclude that DomainDLRS is the preferable alternative.

Also on biological data, DomainDLRS clearly outperforms than MrBayesMPR by providing more robust (gene and) domain trees (i.e., with better posterior clade probabilities), clearer and more convincing predictions of domain evolution (both in terms of the number and the position of duplication loss events), and better reconstructions of ancestral genes in terms of domain content.

In particular, our framework gives a reliable analysis of the recent evolution, i.e., of the events succeeding the root of the gene tree, of the zinc-finger families we consider. From our analysis, it appear that single and segmental duplications of zinc-finger domains are more or less equally frequent and, hence, that more domains are affected by segmental duplications.

We make a note that the reconciliation of ancient evolution, that is, events occurring prior the root of the gene tree, is an hard problem, regardless of method; its solution rather lies in breaking up the root edge by additional sampling to provide more constraints for the reconciliation. However, it is also well-known that with erroneous tree-reconstruction, reconciliations will tend to predict an over-abundance of ancient evolutionary events. It is clear from our results that, while this is a major problem woth MrBAyes+MPR, it is nearly non-existant from DomainDLRS.

It has earlier been observed that, when applied to domains, the posterior distribution inferred by MrBayes typically is too flat to allow proper conclusions [15] and we show that DomainDLRS solves this problem. DomainDLRS clade support values display a distribution much more centered around higher support values compared to MrBayesMPR.

Our framework does not take advantage of the order in which the domains occur in the genes, which make it possible to evaluate performance on biological data based on a parsimony criterion for domain reordering across the ancestral vertices. DomainDLRS provides a much more parsimonious reconstruction of domain order, with fewer and more reasonable changes in domain order, than MrBayesMPR.

Our results clearly show that it is possible and in many cases preferable to analyse the evolution of domains and evade aligning genes when analysing gene families that have been exposed to domain duplications. Our model does not include domain swapping events, i.e., transfer of a domain from one gene to another contemporary gene, which would render it possible to analyse a larger set of multi domain proteins. Although it is easy to formulate a model that it includes domain swapping, e.g., following models for lateral gene transfer [25], devising an efficient associated inference framework currently constitutes a substantial challenge.

## 6 Methods

### 6.1 DomainDLRS

Each hyper-parameter in *θ*, i.e., duplication-loss rates and mean, variance of the rate model, is assigned a uniform prior over [0, *α*], where *α* is sufficiently high. For *δ* and *μ* we used *α* = 10 and for m and *v* we used *α* = 1000. We here used the same substitution model, i.e. JC69, for all domains in the analyses of biological and synthetic dataset.

The DomainDLRs software has not yet included the automatized convergence testing to estimate the number of samples (iterations) needed for convergence. We tested the convergence of DomainDLRS by running the pilot run on biological datasets. We ran DomainDLRS mcmc chains for 1 million iterations with every 100th iterations sampled. In case of large domain families such as ZNF91, we considered 2 million iteration to be sufficient for thier convergence.

For convergence analysis, we used R software package CODA [26]. To test that our unnormalized model density has been converged we used Heidelberger and Welch’s convergence diagnostic from coda package.

The Heidelberger and Welch’s convergence test is applied on resulting single chain from DomainDLRS to test its stationarity. The null hypothesis that the sampled values are drawn from a stationary distribution, is tested by Cramer-von-Mises statistic. The test has been applied successively first to whole chain, then after disacrding first 10%, 20%,…, of the chain until either the null hypothesis is accepted for given level of significance, or 50% of the chain has been discarded and failed to pass the stationarity. More detail can be found web page of CODA pacakge [26].

In all most all of the Zinc-Finger families pilot mcmc run, the chain has passed the Heidelberger and Welch’s convergence test with 95% significance test. The traces of unnormalized model densities are given in mcmc_post_analysis folder (as part of supplementary material).

### 6.2 MrBayesMPR

The MrBayesMPR method formalizes the standard approach of independent reconstruction of domain trees, followed by computation of MPR between domain tree and a given gene tree. Briefly, MrBayes is used to estimate a domain tree from the domain multialignment. For the MPR, we then use Notung (version 2.6) [27], which reconcile the domain tree to the genetree and simultaneouly root domain tree to minimize duplication and loss events. We also use the MrBayesMPR method to reconcile gene trees with the species tree.

### 6.3 Synthetic data generation

We generated three sets of synthetic gene families under the DomainDLRS model, each containing 100 synthetic gene families, here referred to as, SZNF91, SZNF468, and SZNF679, based on the MAP parameters obtained in the DomainDLRS analyses of the gene families ZNF91, ZNF468, and ZNF679, respectively, see Section 6.4. The parameters are described in Supplementary Table **??**. For the synthetic data, the GenPhyloData software package [28] was used to generate dated and length equipped trees; the Seq-Gen program, from [29], was used to generated the sequence data from the length equipped domain trees. The species tree from Nowick et. al. [30] consisting of 4 species, i.e., Human, Chimpanzee, Orangutan, and Rhesus Macaque, was used in the in the generation, as well as the subsequent analysis. In the case of the mono-copy gene family ZNF91, for which the gene is likely to coincide with the species tree, we also used the species tree as gene tree when generating the domain trees for SZNF91.

The output from each generation comprises a gene tree, a domain tree, reconciliations between the species tree and the gene tree gene and between the domain tree and the gene tree, respectively, and a multiple sequence alignment of the domain sequences. To enable a comparison of the accuracy of our method’s gene tree inference with that of MrBayesMPR, also gene alignments were generated. For this purpose, domain sequences were concatenated according to a random model as well as a tandem model. In the random model, gene alignment data were constructed by concatenating zinc finger domains placed in random order; notice though that a single KRAB domain was always placed first, in agreement with the biological datasets. In the tandem model, the zinc finger domains were ordered according to a post-order traversal of the respective domain tree.

### 6.4 Biological Data

We applied DomainDLRS and MrBayes-MPR to experimental data from the following families, ZNF91, ZNF558/557, ZNF468/28, ZNF679/716, ZNF611/600, and ZNF764/747. The sequence data was obtained from publications or downloaded from the databases NCBI, Ensembl, and the KRAB associated Zinc-Finger catalogue [31], the latter available at http://znf.igb.uiuc.edu/ (for accession details, see Supplementary Table S2). The domain coordinates were identified based on protein sequence, using the SMART domain profile database [32], and then used to extract the DNA sequences of domains for our analysis. The multiple sequence alignment program Muscle [33] was used, with default parameter setting, to align the domain sequences. Muscle was also used to align gene sequences for the MrBayesMPR gene tree reconstruction; for gene sequences, sequence regions not belonging to a domain were removed. For of each domain family, the source, the gene identifiers, and the domain information are specified in SuppementaryTable **??**.

### 6.5 Reconciliation heatmaps

A reconciliation heat map is an visualization of a reconciled MAP guest tree, inside the host tree, with a color coding of edges vertices indicating the posterior probability of the corresponding clades. All reconciliation heatmaps were created using an in-house extension of the PrIMETV software [34], that uses a parsimony-based algorithm to reconstruct domain order at internal gene tree vertices. The algorithm algorithm greedily orders domain vertices to minimize domain rearrangements, and is not guaranteed to give the optimal solution, but rather a rough estimate of domain shuflling event.

## References

[1] Örjan Åkerborg, Bengt Sennblad, Lars Arvestad, and Jens Lagergren. Simultaneous bayesian gene tree reconstruction and reconciliation analysis. Proceedings of the National Academy of Sciences, 106(14):5714–5719, 2009.

[2] Matthew D Rasmussen and Manolis Kellis. A bayesian approach for fast and accurate gene tree reconstruction. Molecular Biology and Evolution, 28(1):273–290, 2011.

[3] Bastien Boussau, Gergely J Szöllősi, Laurent Duret, Manolo Gouy, Eric Tannier, and Vincent Daubin. Genome-scale coestimation of species and gene trees. Genome research, 23(2):323–330, 2013.

[4] Peter L Oliver, Leo Goodstadt, Joshua J Bayes, Zoë Birtle, Kevin C Roach, Nitin Phadnis, Scott A Beatson, Gerton Lunter, Harmit S Malik, and Chris P Ponting. Accelerated evolution of the prdm9 speciation gene across diverse metazoan taxa. PLoS Genet, 5(12):e1000753, 2009.

[5] Frank MJ Jacobs, David Greenberg, Ngan Nguyen, Maximilian Haeussler, Adam D Ewing, Sol Katzman, Benedict Paten, Sofie R Salama, and David Haussler. An evolutionary arms race between krab zinc-finger genes znf91/93 and sva/l1 retrotransposons. Nature, 516(7530):242–245, 2014.

[6] Åsa K Björklund, Diana Ekman, and Arne Elofsson. Expansion of protein domain repeats. PLoS Comput Biol, 2(8):e114, 2006.

[7] Jung-Hoon Han, Sarah Batey, Adrian A Nickson, Sarah A Teichmann, and Jane Clarke. The folding and evolution of multidomain proteins. Nature Reviews Molecular Cell Biology, 8(4):319–330, 2007.

[8] Hamsa Dhwani Tadepally and Muriel Aubry. Evolution of c2h2 zinc-finger gene families in mammals. eLS.

[9] AT Hamilton, S Huntley, J Kim, E Branscomb, and L Stubbs. Lineage-specific expansion of krab zinc-finger transcription factor genes: implications for the evolution of vertebrate regulatory networks. In Cold Spring Harbor symposia on quantitative biology, volume 68, pages 131–140. Cold Spring Harbor Laboratory Press, 2003.

[10] Katja Nowick, Tim Gernat, Eivind Almaas, and Lisa Stubbs. Differences in human and chimpanzee gene expression patterns define an evolving network of transcription factors in brain. Proceedings of the National Academy of Sciences, 106(52):22358–22363, 2009.

[11] Gernot Wolf, David Greenberg, and Todd S Macfarlan. Spotting the enemy within: Targeted silencing of foreign dna in mammalian genomes by the krüppel-associated box zinc finger protein family. Mobile DNA, 6(1):1, 2015.

[12] James H Thomas and Sean Schneider. Coevolution of retroelements and tandem zinc finger genes. Genome research, 21(11):1800–1812, 2011.

[13] Morris Goodman, John Czelusniak, G William Moore, AE Romero-Herrera, and Genji Matsuda. Fitting the gene lineage into its species lineage, a parsimony strategy illustrated by cladograms constructed from globin sequences. Systematic Biology, 28(2):132–163, 1979.

[14] Ali Tofigh, Michael Hallett, and Jens Lagergren. Simultaneous identification of duplications and lateral gene transfers. IEEE/ACM Transactions on Computational Biology and Bioinformatics (TCBB), 8(2):517–535, 2011.

[15] Maureen Stolzer, Katherine Siewert, Han Lai, Minli Xu, and Dannie Durand. Event inference in multidomain families with phylogenetic reconciliation. BMC bioinformatics, 16(Suppl 14):S8, 2015.

[16] Lars Arvestad, Ann-Charlotte Berglund, Jens Lagergren, and Bengt Sennblad. Gene tree reconstruction and orthology analysis based on an integrated model for duplications and sequence evolution. In Proceedings of the eighth annual international conference on Resaerch in computational molecular biology, pages 326–335. ACM, 2004.

[17] Joel Sjöstrand, Ali Tofigh, Vincent Daubin, Lars Arvestad, Bengt Sennblad, and Jens Lagergren. A Bayesian Method for Analyzing Lateral Gene Transfer. Systematic Biology, 63(3):409–420, March 2014.

[18] Bengt Sennblad and Jens Lagergren. Probabilistic orthology analysis. Systematic biology, 58(4):411–424, 2009.

[19] Ikram Ullah, Joel Sjöstr and, Peter Andersson, Bengt Sennblad, and Jens Lagergren. Integrating sequence evolution into probabilistic orthology analysis. Systematic biology, page syv044, 2015.

[20] Lars Arvestad, Ann-Charlotte Berglund, Jens Lagergren, and Bengt Sennblad. Bayesian gene/species tree reconciliation and orthology analysis using mcmc. Bioinformatics, 19(suppl 1):i7–i15, 2003.

[21] Mark A Beaumont. Estimation of population growth or decline in genetically monitored populations. Genetics, 164(3):1139–1160, 2003.

[22] Christophe Andrieu and Gareth O Roberts. The pseudo-marginal approach for efficient monte carlo computations. The Annals of Statistics, pages 697–725, 2009.

[23] Joseph Felsenstein. Evolutionary trees from dna sequences: a maximum likelihood approach. Journal of molecular evolution, 17(6):368–376, 1981.

[24] Matthew W Hahn. Bias in phylogenetic tree reconciliation methods: implications for vertebrate genome evolution. Genome biology, 8(7):1–9, 2007.

[25] Joel Sjöstrand, Ali Tofigh, Vincent Daubin, Lars Arvestad, Bengt Sennblad, and Jens Lagergren. A Bayesian Method for Analyzing Lateral Gene Transfer. Syst Biol, 63(3):syu007–420, February 2014.

[26] Martyn Plummer, Nicky Best, Kate Cowles, and Karen Vines. Coda: convergence diagnosis and output analysis for mcmc. R news, 6(1):7–11, 2006.

[27] Kevin Chen, Dannie Durand, and Martin Farach-Colton. NOTUNG: A Program for Dating Gene Duplications and Optimizing Gene Family Trees. J. Comput. Biol., 7(3-4):429–447, August 2000.

[28] Joel Sjöstrand, Lars Arvestad, Jens Lagergren, and Bengt Sennblad. Genphylodata: realistic simulation of gene family evolution. BMC bioinformatics, 14(1):209, 2013.

[29] Andrew Rambaut and Nicholas C Grass. Seq-gen: an application for the monte carlo simulation of dna sequence evolution along phylogenetic trees. Computer applications in the biosciences: CABIOS, 13(3):235–238, 1997.

[30] Katja Nowick, Aaron T Hamilton, Huimin Zhang, and Lisa Stubbs. Rapid sequence and expression divergence suggest selection for novel function in primate-specific krab-znf genes. Molecular biology and evolution, 27(11):2606–2617, 2010.

[31] Stuart Huntley, Daniel M Baggott, Aaron T Hamilton, Mary Tran-Gyamfi, Shan Yang, Joomyeong Kim, Laurie Gordon, Elbert Branscomb, and Lisa Stubbs. A comprehensive catalog of human krab-associated zinc finger genes: insights into the evolutionary history of a large family of transcriptional repressors. Genome research, 16(5):669–677, 2006.

[32] Jörg Schultz, Frank Milpetz, Peer Bork, and Chris P Ponting. Smart, a simple modular architecture research tool: identification of signaling domains. Proceedings of the National Academy of Sciences, 95(11):5857–5864, 1998.

[33] Robert C Edgar. Muscle: multiple sequence alignment with high accuracy and high throughput. Nucleic acids research, 32(5):1792–1797, 2004.

[34] Bengt Sennblad, Eva Schreil, Ann-Charlotte Berglund Sonnhammer, Jens Lagergren, and Lars Arvestad. Primetv: a viewer for reconciled trees. BMC bioinformatics, 8(1):148, 2007.

